# De-novo generation of novel phenotypically active molecules for Chagas disease from biological signatures using AI-driven generative chemistry

**DOI:** 10.1101/2021.12.10.472084

**Authors:** Michal Pikusa, Olivier René, Sarah Williams, Yen-Liang Chen, Eric Martin, William J. Godinez, Srinivasa P S Rao, W. Armand Guiguemde, Florian Nigsch

## Abstract

Designing novel molecules with targeted biological activities and optimized physicochemical properties is a challenging endeavor in drug discovery. Recent developments in artificial intelligence have enhanced the early steps of *de novo* drug design and compound optimization. Herein, we present a generative adversarial network trained to design new chemical matter that satisfies a given biological signature. Our model, called pqsar2cpd, is based on the activity of compounds across multiple assays obtained via pQSAR (profile-quantitative structure–activity relationships). We applied pqsar2cpd to Chagas disease and designed a novel molecule that was experimentally confirmed to inhibit growth of parasites *in vitro* at low micromolar concentrations. Altogether, this approach bridges chemistry and biology into one single framework for the design of novel molecules with promising biological activity.

## Introduction

Recently, artificial intelligence has been successfully applied in early drug discovery settings to generate novel compounds and optimize existing ones towards better physicochemical properties. Machine learning models can learn the association of molecular representations with molecular properties^1^, and generative models were proposed to help with the creation of new chemical matter based on historical experimental data^2^. Several methods have been developed based on techniques such as Reinforcement Learning^3,4,5^, Variational Auto Encoders (VAE)^6,7^, and Generative Adversarial Networks (GAN)^8,9^.

While most approaches require the structure of a seed compound for generating novel structures, several studies reported the *in silico* generation of chemical matter from transcriptomic data^10^ and protein sequences^11^. As opposed to a seed molecule, these methods start with an expression profile and protein sequence, respectively. We built upon these latter approaches to present a solution for generation of new molecular structures based on a given biological activity profile of interest, more specifically EC_50_ activities against selected targets. The uniqueness of our approach is that a bioactivity profile, or “fingerprint,” can be turned into a novel compound capable of inducing a similar activity profile. In this report, we leveraged a machine learning method, pQSAR (profile Quantitative Structure-Activity Relationships)^12^, to generate a bioactivity profile. pQSAR is an automated, massively multitask, QSAR method that predicts biological activities, and has accurate models for more than 8500 Novartis assays, spanning 51 annotated target sub-classes, as well as over 4000 phenotypic assays. Our approach called *pqsar2cpd* relies on a pQSAR-based fingerprint of biological activities and can produce completely novel and unbiased scaffolds, which gives an additional advantage over chemical seed-based models.

Artificial intelligence is poised to speed the process of drug discovery in various diseases such as Chagas disease, an infectious disease which was selected as a test case for identifying new potential small molecule inhibitors via generative chemistry. Chagas disease is a neglected parasitic disease affecting around 6 million people in South America alone, and there is an urgent need for new treatments^13^. Hence, the rationale for investigating and leveraging artificial intelligence is not only to speed up the drug discovery process, but also to minimize costs. In this report, we highlight how pqsar2cpd was applied to Chagas disease and proposed a compound that was confirmed experimentally *in vitro* with antiparasitic activity.

Although various generative chemistry models based on deep learning have been published with proven *in silico* success, only a handful have provided experimental *in vitro* confirmation after chemical synthesis. This constitutes a considerable obstacle in connecting generative models and medicinal chemistry^3^. To our knowledge, this is the first report to demonstrate successful *in vitro* validation of compounds generated from a biological activity profile using Generative Adversarial Networks.

## Results

### pqsar2cpd

pqsar2cpd is based on Generative Adversarial Networks^14^. GANs are deep neural networks trained to generate synthetic data that mirrors the feature distribution of the training data. They consist of a generator, which captures the training data distribution and produces synthetic samples that closely follow it, and a discriminator that estimates the probability of a sample coming from the generator rather than from the training data. The end goal of the training of such networks is to have a generator produce synthetic samples that the discriminator cannot distinguish from training samples.

In this work, we used a conditional GAN (c-GAN) that includes the pQSAR bioactivity fingerprints of the training compounds for conditioning the generator. In comparison to an unconditioned GAN, a GAN conditioned on biological fingerprints aims to produce not only novel chemical structures but rather the structures that are likely to have a given activity profile. c-GANs therefore require additional input to the generator, in this case pQSAR fingerprints, as well as a conditional neural network to assess the probability of a generated structure corresponding to a given pQSAR fingerprint **(Figure 1a-1b)**.

**Figure 1a.**
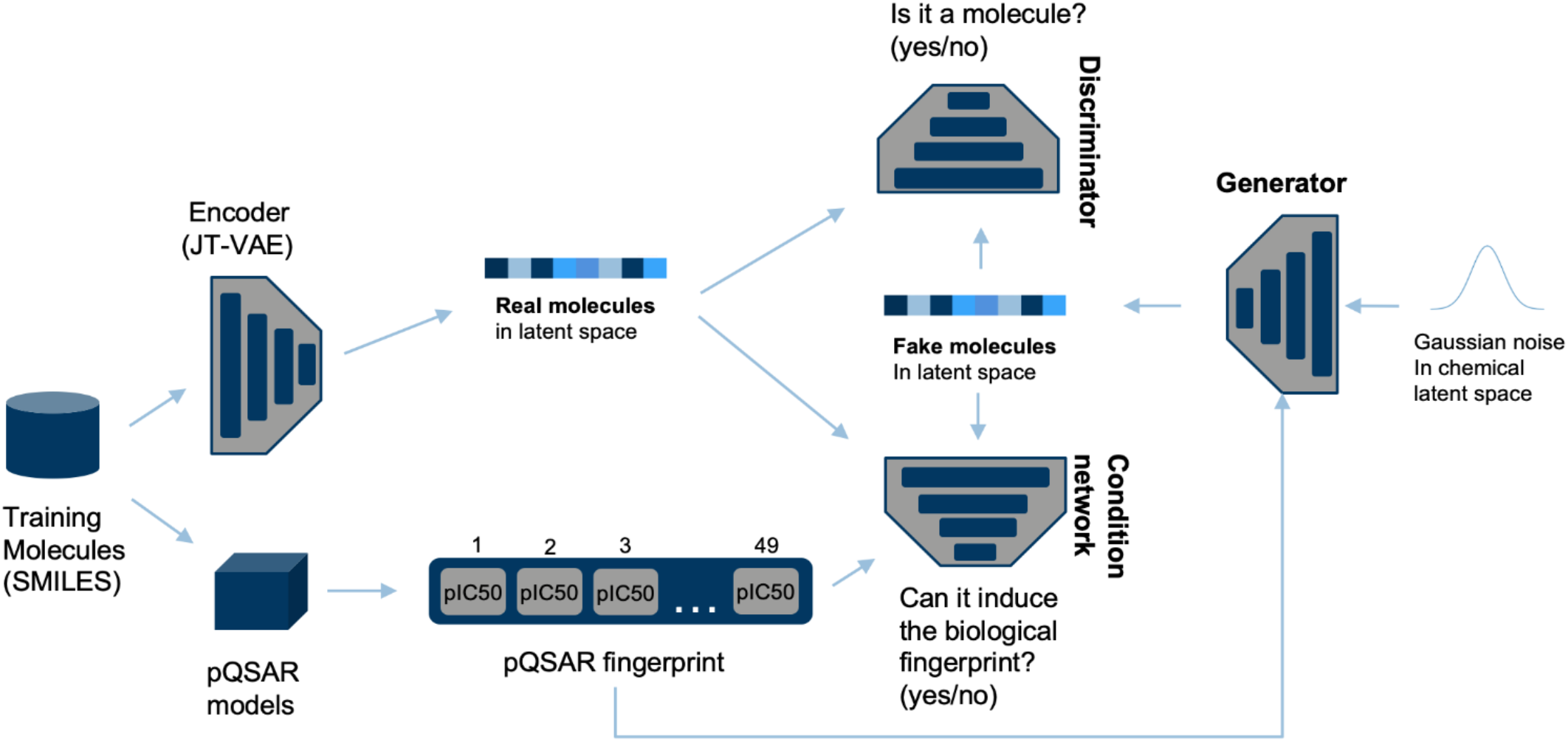
Diagram of pqsar2cpd in the training phase

**Figure 1b.**
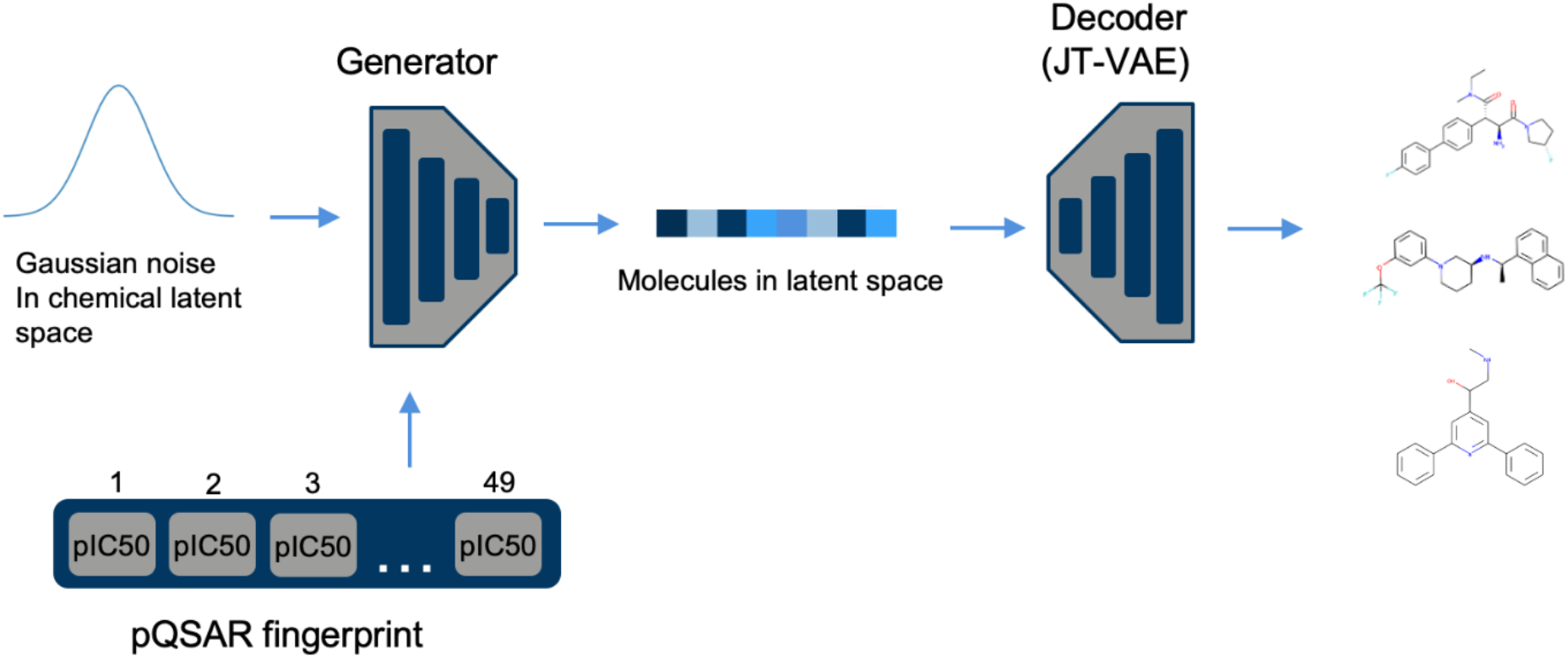
Diagram of pqsar2cpd in the inference phase

We selected a subset of the thousands of assays covered by pQSAR in order to create a more specific antiparasitic fingerprint for Chagas. A total of 49 assays were selected that are representative of, or related to, the involved parasite: *Tryponasoma cruzi, Tryponasoma brucei, Leishmania donovani*, and *Plasmodium falciparum*. All these are kinetoplastid parasites, except *Plasmodium falciparum*, and share significant genome similarity. The assays used for fingerprinting were mainly cellular assays determining the growth inhibitory potential of compounds.

The representation of molecular structures *in silico* is a central problem of cheminformatics and paramount for generative chemistry. Whereas encoding of discrete chemical structures into a continuous latent space is always feasible, not every point in that latent space corresponds to a physically possible molecule.^17^ The latent space for our modelling is based on a method that is guaranteed to decode only into chemically valid structures. More specifically, we used a Junction-Tree Variational Autoencoder (JT-VAE)^16^ that was trained to represent anti-parasitic compounds as implemented in JAEGER^15^ (see Methods). pqsar2cpd was trained on a set of 16,000 latent compound vectors coming from JAEGER along with corresponding pQSAR activity fingerprints across the 49 selected anti-parasitic assays (see Methods).

### Molecule generation for Chagas and *in silico* model evaluation

Our goal was to generate structurally novel growth inhibitors of *T. cruzi* that mimic the pQSAR profiles of eight known compounds active against *T. cruzi in vitro*. Thus, these eight pQSAR profiles (across the same set of 49 anti-parasitic assays that were used for training) were used as seed profiles for pqsar2cpd. This resulted in 100 chemically valid molecules out of which 88 were unique (with 56 unique scaffolds, based on Murcko scaffold extraction).

Any molecule proposed by pqsar2cpd needs to meet a high standard: it should be novel; it should be synthetically tractable; it should have reasonable medicinal chemistry properties; and its pQSAR fingerprint should be similar to the model input.

The 88 generated compounds had a mean structural similarity of 0.16 and 0.19 (Tanimoto coefficient using Morgan fingerprints) to the eight known *T. cruzi* inhibitors and the 16,000 compounds in the training set, respectively (**Figure 2a**). This confirms the generation of novel molecules and the absence of overt copying of the training set. The median synthetic accessibility (SA)^18^ score for the generated molecules was 2.5 (all molecules were below 4.5, a common threshold for “ease of synthesis” used in previous studies^10^), whereas the quantitative estimation of drug-likeness (QED)^19^ value was above 0.6 for 54 (60%) of the molecules (**Figure 2b**). The pQSAR profiles of the generated compounds showed a mean cosine similarity of 0.32 to the seed profiles and 0.14 to the training profiles. A pQSAR model specific to *T. cruzi* predicted a pIC_50_ greater than 4.5 (i.e., less than 20 μM) for 81 (92%) of the molecules (**Figure 2c**). Additional search for close analogs in 3 public repositories, i.e. PubChem, Zinc, and eMolecules, containing over 170 million molecules, revealed no molecules with structures identical to generated ones, thus providing additional confirmation of their novelty. The median Tanimoto similarity of generated compounds to the closest existing compounds was 0.64 (**Figure 2d**), while the median similarity to the only two FDA-approved drugs for Chagas disease, Benznidazole and Nifurtimox, was 0.18 and 0.09 respectively (**Figure 2e**).

**Figure 2a.**
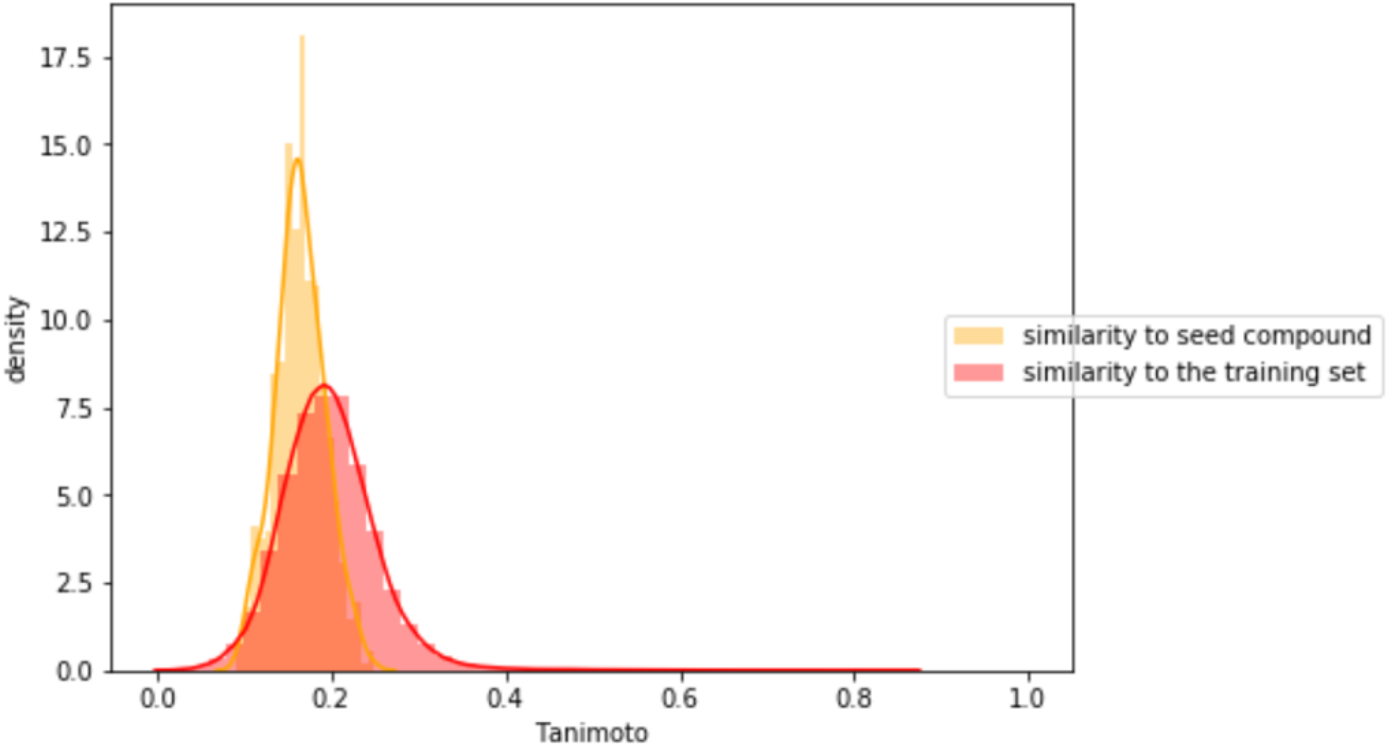
Distribution of Tanimoto similarities of generated compounds (n=88) to training (n=16,000) and seed molecules (n=8)

**Figure 2b.**
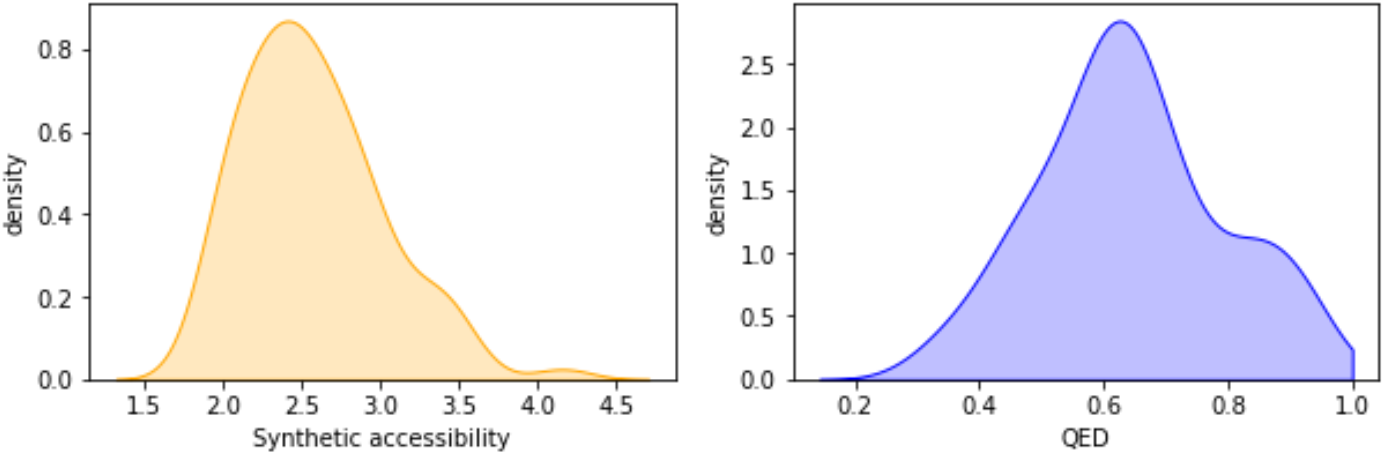
Distribution of molecular properties of generated compounds (n=88)

**Figure 2c.**
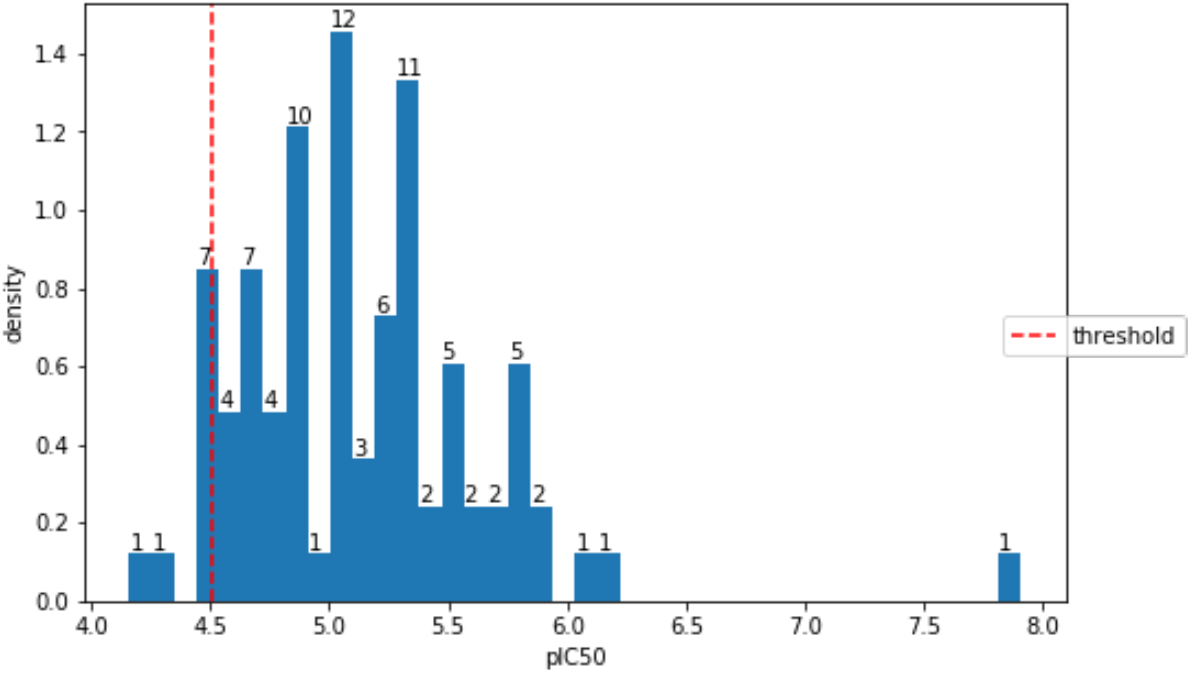
Distribution of predicted pIC50 activities (n=88) on T. cruzi assay

**Figure 2d.**
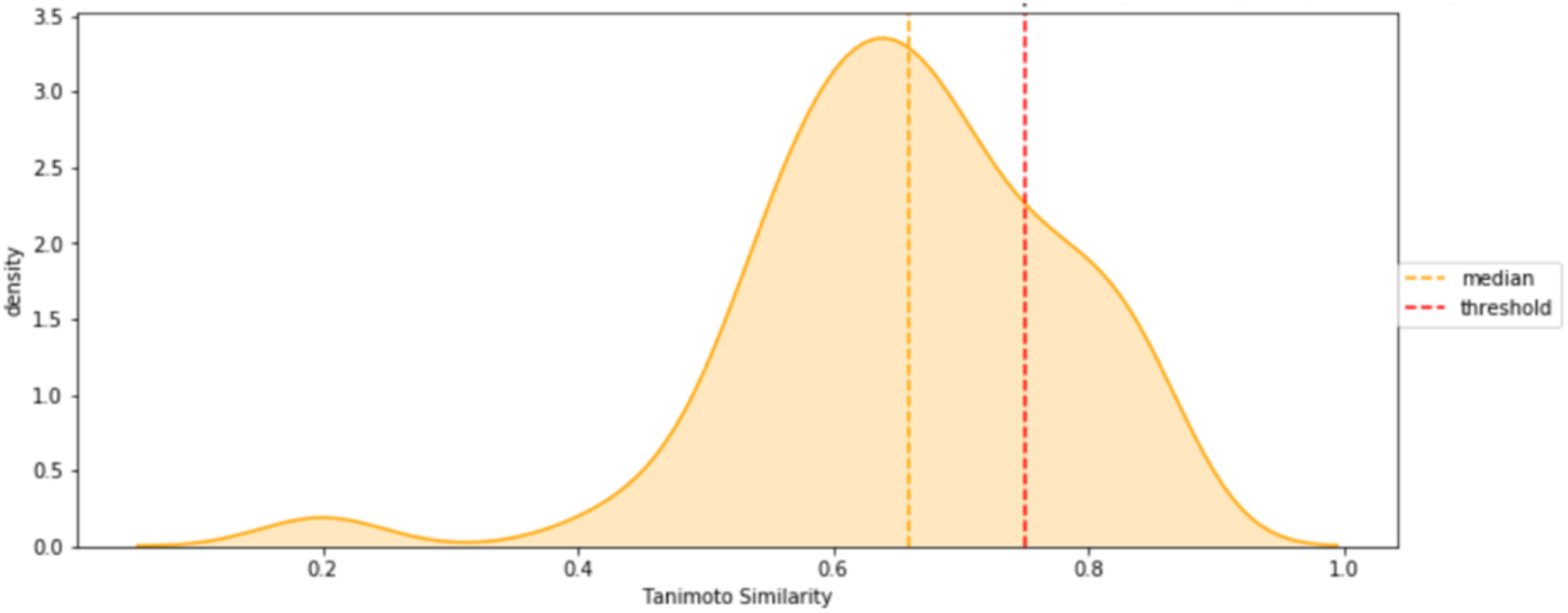
Distribution of similarities of generated compounds (n=88) to the closest existing compound in public databases, PubChem, Zinc, and eMolecules (n ~ 178M)

**Figure 2e.**
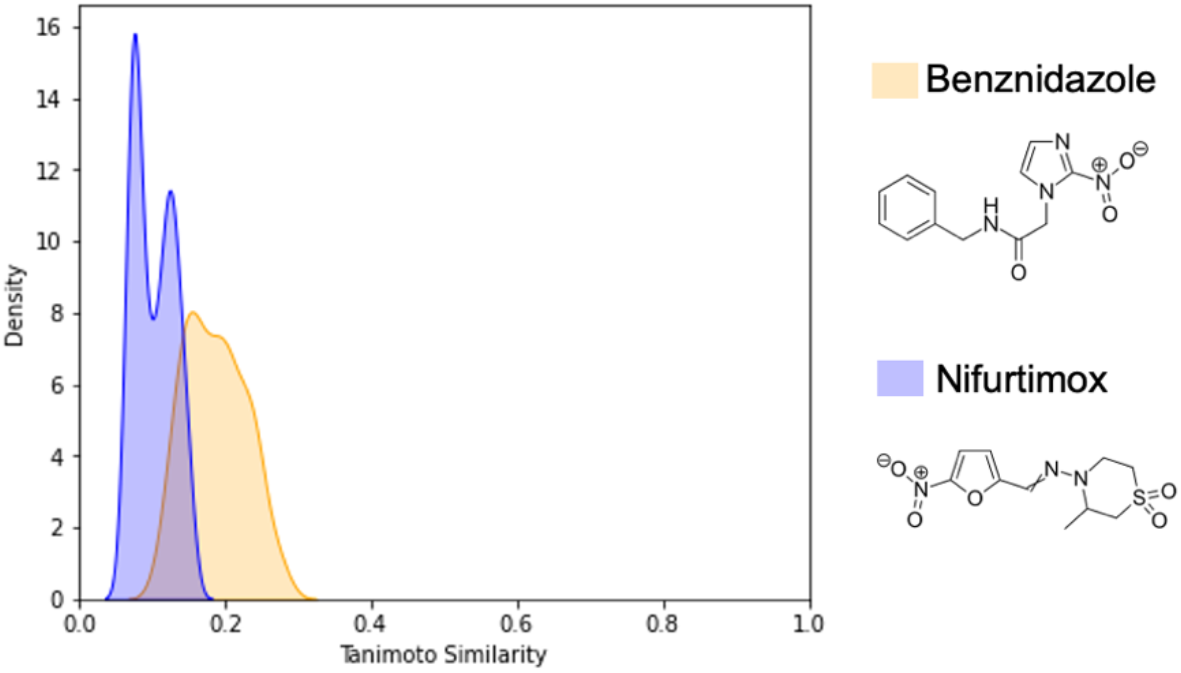
Distribution of similarities of generated compounds (n=88) to FDA-approved drugs for Chagas disease (n=2)

Taken together, these data confirmed that our model was performing as intended: the generated molecules were sufficiently distinct in structure from known compounds, while most of them fell into the chemical space that was (likely) amenable to synthesis as well as medicinal chemistry efforts to optimize ADMET properties. Paramount though to our method’s purpose is the generation of molecules according to a predefined biological fingerprint. Our assessment showed that the activity profiles of the generated molecules were indeed more similar to the seeds than to the training profiles. Follow-up with a specific *T. cruzi* model further confirmed the *in silico* validity of the method.

### Selection of compounds for *in vitro* testing and experimental results

We prioritized the best compounds for synthesis and *in vitro* testing based on pQSAR predicted activity against *T. cruzi* and chemistry assessment. A final set of three compounds with predicted pIC_50_ values of 7.88, 6.4, and 6.4 were chosen (**Figure 3a**). Compounds **GEN-1** and **GEN-2** were successfully synthesized and compound **GEN-3**, with the highest predicted pIC_50_, was ultimately determined to be synthetically intractable and was deprioritized. Compounds **GEN-1** and **GEN-2** were tested *in vitro* against *T. cruzi* grown in 3T3 mice-derived fibroblast host cells. Upon compound addition and incubation over 6 days, **GEN-1** was moderately active with an EC_50_ value of 4 μM whereas **GEN-2’s** EC_50_ was greater than 20 μM. In comparison, the approved drug benznidazole used as control in this assay was 5-fold more potent with an EC_50_ of 0.8 μM. Lastly, we investigated potential off-targets associated with cytotoxicity. **GEN-1** and **GEN-2** were tested against 3T3 host cells to investigate whether *T. cruzi* activity was driven by host cell killing. After incubation over 4 days, **GEN-1** and **GEN-2** displayed very low levels of cytotoxicity with EC_50_ values greater than 20 μM for both, whereas the control compound puromycin was highly active with an EC_50_ of 1.4 μM (**Figure 3b**).

**Figure 3a.**
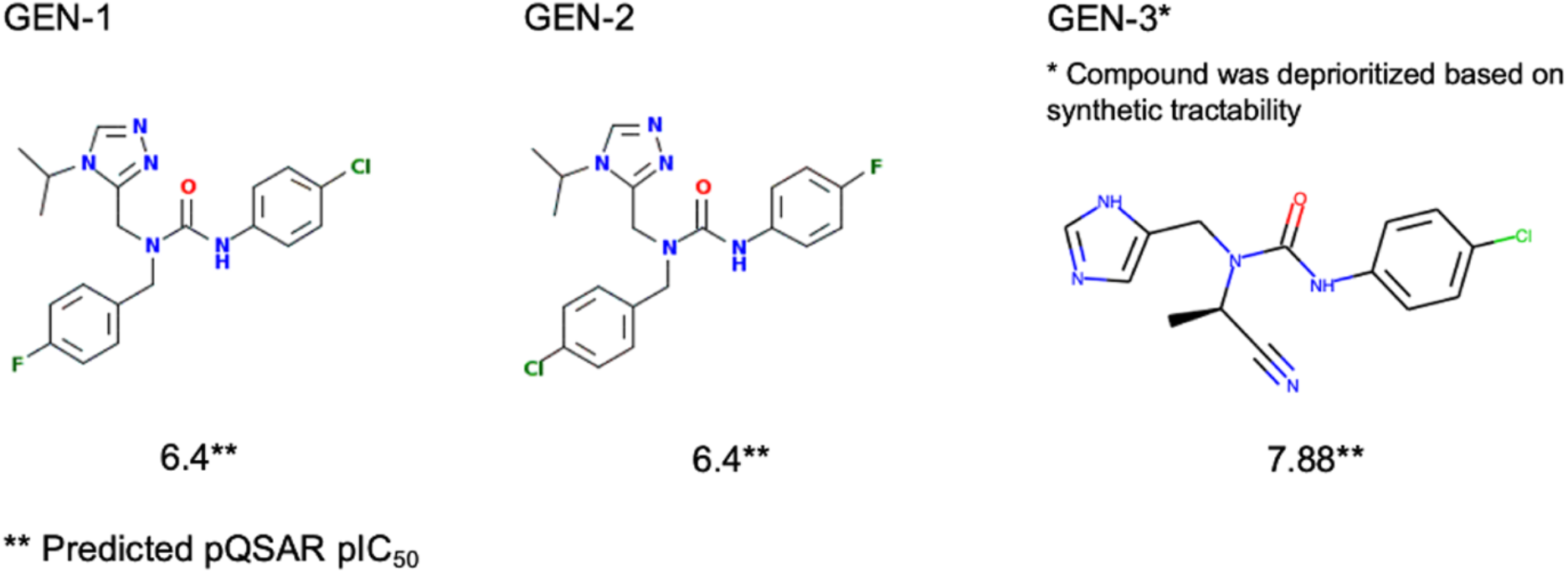
Generated compounds prioritized for synthesis

**Figure 3b.**
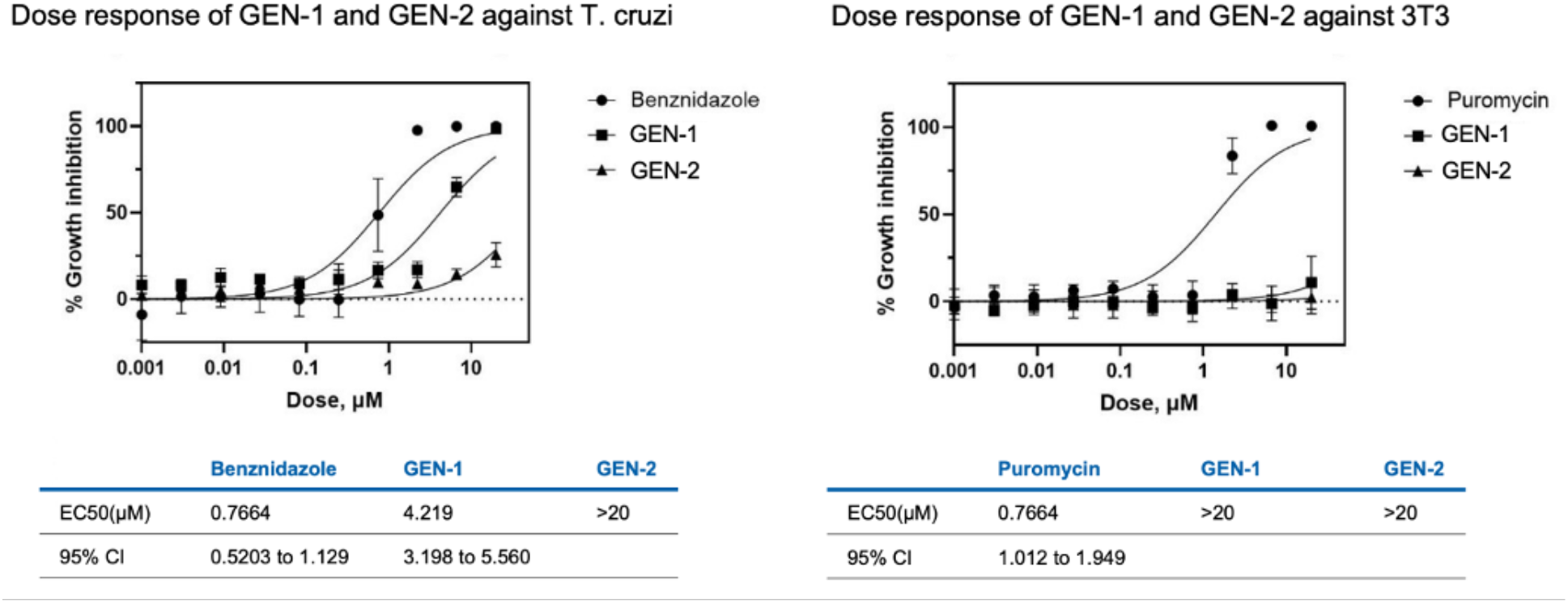
Dose-response results of synthesized compounds against T. cruzi and 3T3

## Discussion

In this study, we reported pqsar2cpd, a deep learning method based on a conditional generative adversarial network to generate novel molecules fitting a defined bioactivity profile (pIC_50_ fingerprint) from pQSAR multitask models. Pqsar2cpd is a generative chemistry approach to directly translate multi-parametric biological fingerprints into chemical matter, inspired by previous studies that used gene expression signatures to the same end^10^. pQSAR is a published method that predicts accurate pIC_50_ profiles for thousands of biochemical and phenotypic assays. Although our models are built on proprietary screening data, a custom pqsar2cpd model can be trained on publicly available data such as ChEMBL. In contrast to previous studies that generated <10% chemically valid structures^10^, we used a graph-based encoding/decoding scheme for molecular structures that is guaranteed to result in only chemically valid molecules in a predefined anti-parasitic chemical space. Moreover, the molecules sampled from this space had low structural similarity to the training set, while retaining the desired biological activity profile, as well as drug likeness and synthetic accessibility. The generation of dissimilar molecules from the training set was indicative of a propensity for scaffold hopping, allowing the model to jump to new areas of chemical space through combinations of the learned fragments in novel yet chemically feasible ways. In addition to extensive computational validation, we used pqsar2cpd in our Chagas drug discovery efforts to design a novel molecule with confirmed *in vitro* activity against *T. cruzi* parasites. Despite this successful combination of computational, chemical and biological expertise, further work is needed to shed more light on important aspects: for example, compound **GEN-2** was not active *in vitro*, despite being a close analog of **GEN-1**. In hindsight, the prioritization *via* predicted pIC_50_ values was suboptimal in this case due to the indistinguishable computational representation (Morgan fingerprints) of the two input molecules to the employed pQSAR model. Another important factor is the discrepancy of the predicted synthetic tractability of **GEN-3** to actual synthetic chemistry efforts required, resulting in the abandonment after multiple failed attempts. We are actively using and refining pqsar2cpd in ongoing drug discovery projects in order to identify ideal use cases and concomitant limitations and to increase the biological relevance of the proposed molecules along with their attractiveness for medicinal chemistry.

## Methods

### pQSAR

Profile-quantitative structure-activity relationship (pQSAR)^12^ is a two-level, stacked, machine-learning method. In level one, a profile of conventional, single-assay random forest regression (RFR) models were trained on 13,500 biochemical and cellular pIC_50_ assays using Morgan 2 substructural fingerprints as compound descriptors. In level two, a panel of partial least squares (PLS) regression models were built for the same 13,500 assays, now using the profile of RFR pIC_50_ predictions as compound descriptors. This 2-level process transfers learning between the assays, dramatically increasing the accuracy of the models. The median correlation between the pIC_50_ predictions and experimental values on compounds very unlike the training sets was 0.52, comparable to 4-concentration experimental IC_50_s. The model was used to predict pIC_50_ values for both training and seed compounds used in the training, and the predictions of the GAN.

### pqsar2cpd architecture

#### Molecule encoder/decoder - JAEGER/JT-VAE

JT-VAE represents molecules through both a junction tree subsuming the topology of substructures within a molecule, as well as through a molecular graph capturing the atom and bond structure of a molecule. Passing both through a set of neural networks, it can map molecules to a latent space representation, which in turn can be used to predict a junction tree back and reconstruct a molecular graph of the input molecule.

The goal during training of JAEGER is to improve the performance of the JT-VAE model in terms of its ability to 1) reconstruct accurately both trees and graphs, 2) obtain a good approximation of the variational tree and graph posteriors and 3) properly predict pIC_50_s given a latent representation. Hence, JAEGER can validate the quality of the latent space created by JT-VAE.

To train JAEGER and the underlying JT-VAE models we used a set of 16,000 molecules along with pIC_50_ values coming from an internal *Buckner infectious T. Cruzi* assay. Molecules were represented as SMILES, and after training JAEGER, were encoded in a single latent vector of 56-dimensions for each molecule.

With JAEGER trained and validated, the JT-VAE molecule encoder was used to encode molecules into their 56-dimensional latent space vectors needed to train pqsar2cpd, and the decoder was used to decode compounds generated by it after training back into SMILES. Hence, pqsar2cpd operates in an established latent space, and the quality of generated molecules depends on its quality. Since previous papers reported a low rate of valid molecules generated using GANs with SMILES-based molecular encoders^10^, using JT-VAE with its 100% validity scores for encoding and decoding ensures that our GAN is working in a valid latent space to begin with.

#### GAN - Generator

The generator receives two inputs: a 56-dimensional noise vector sampled from a normal distribution, and the condition vector consisting of a pQSAR fingerprint of 49 z-scored pIC_50_ values. The two inputs are processed separately with a 2-layer multilayer perceptron (MLP), with 32 nodes in each layer, followed by a LeakyRelu activation function and a batch normalization procedure. Resulting tensors are then concatenated and passed as input to another 2-layer MLP. The first layer is a 32-node layer followed by a LeakyRelu activation function. The second layer is the output layer with the dimensionality of the latent space (56 dimensions), followed by a Tanh activation function.

#### GAN - Discriminator

As the discriminator assesses the probability of an input being close to the training data distribution, it receives a single input of a 56-dimensional vector representing a molecule in latent space. The neural network of the discriminator consists of a 3-layer MLP. The first two 32-node layers are followed by a LeakyRelu activation function, while the last one with a single node is followed by a sigmoid activation function. To reduce overfitting, a dropout layer with a rate of 0.4 was implemented following the first and the second layers.

#### GAN - Conditional network

In order to calculate the probability of inducing a certain pQSAR profile of activities, a conditional network was employed. This network receives as input a 56-dimensional compound vector in latent space, and a 49-dimensional pQSAR vector. Both inputs are processed separately by an identical 2-layer MLP, with 32 nodes in each layer, followed by a LeakyRelu activation function and a batch normalization layer. Resulting tensors are then concatenated and passed through a *SubMult+NN* function^21^. The output of this function is used as an input in the following 2-layer MLP, with 32 and 1 nodes, followed by LeakyRelu and sigmoid activation functions respectively.

#### GAN

With the above elements in place, the GAN is defined as taking the noise vector (*z*) and pQSAR vector (*y*) as inputs, and outputting probabilities from the discriminator (*D*(*G*(*z*)) and the conditional network (*C*(*G*(*z,y*)). *D* and *C* are optimized with a *binary cross entropy* loss function.

### Training

The conditional generative adversarial network was trained on the set of 16,000 latent compound vectors coming from JAEGER along with corresponding pQSAR vectors of 49 pIC_50_ values. We trained the model for 400 epochs, with a batch size of 128. The weights of the discriminator and conditional network were updated at each training step, using separate half-batches of real and generated samples, while the weights of the generator were updated every 10 steps. Label smoothing and inverting of labels for real and fake samples was also employed, as it has been shown to improve the training quality ^22^. A SGD optimizer (with a learning rate of 0.005) was used for the discriminator, while an Adam optimizer (with a learning rate of 0.005) was used for the generator. The whole architecture was implemented in Tensorflow^23^. It took 4 hours to train the model on a single Tesla K80 GPU with 8GB of memory.

### Compound synthesis

The detailed synthesis protocol of **GEN-1** and **GEN-2** is included in Supplementary materials.

### *T. cruzi* and 3T3 assay description

The detailed *T. cruzi* and 3T3 assay descriptions and protocols are included in Supplementary materials.

## Author contributions

M.P., W.A.G. and F.N. designed, and led the study. M.P. developed and implemented pqsar2cpd. W.A.G., Y.L.C. and S.P.S.R sampled the biological assays. Y.LC. and S.P.S.R conducted the profiling experiments. O.R. selected and led the synthesis of pqsar2cpd proposed molecules. O.R. and S.W. provided feedback on validity, drug like properties and chemical tractability of pqsar2cpd generated molecules. W.J.G. provided feedback and the framework for valid chemical generation using JAEGER. E.J. Martin provided feedback and pQSAR fingerprints. M.P, W.A.G., and F.N. wrote the manuscript. All authors reviewed the manuscript.

## Corresponding authors

Correspondence to M.P., W.A.G., and F.N.

## Data, Material and Code Availability Statement

### pqsar2cpd

The material, data and software described in this article are of confidential nature and subject to usage restrictions, but may be shared upon execution of respective industry standard agreements with Novartis Pharma AG. Researchers interested in access to the material, data and software may contact Michal Pikusa at.

### pQSAR

Code required to build pQSAR models is available at https://github.com/Novartis/pQSAR

## Acknowledgments

Authors would like to acknowledge funding support from Wellcome Trust (219639_Z_19_Z)

## Supplementary materials

### Synthesis of 3-(4-Chlorophenyl)-1-(4-fluorobenzyl)-1-((4-isopropyl-4*H*-1,2,4-triazol-3-yl)methyl)urea GEN-1

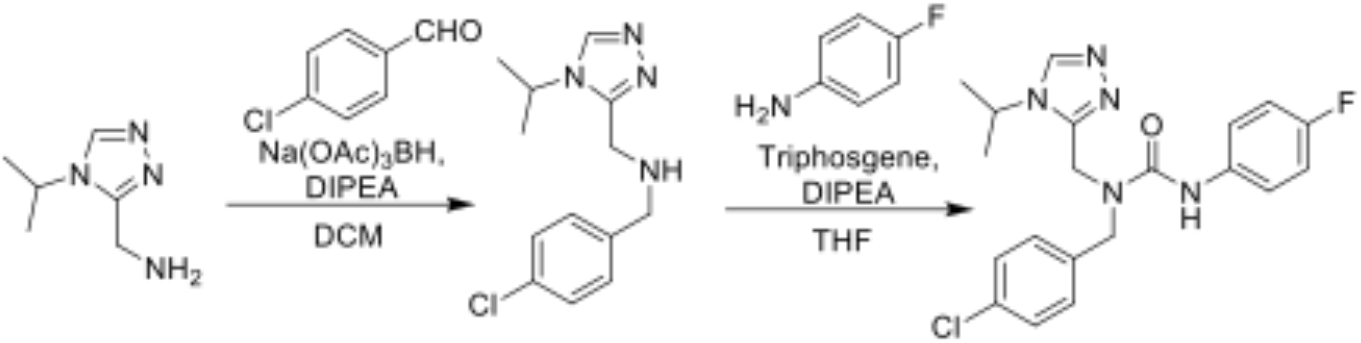

### *N*-(4-Fluorobenzyl)-1-(4-isopropyl-4*H*-1,2,4-triazol-3-yl)methanamine

To a stirred solution of 4-fluorobenzaldehyde (150 mg, 1.21 mmol) in dichloromethane (10 mL) were added (4-isopropyl-4H-1,2,4-triazol-3-yl)methanamine (169 mg, 1.21 mmol), followed by *N,N*-diisopropylethylamine (0.6 mL, 3.63 mmol) and 2 drops of acetic acid. The reaction was cooled to 0 °C, and sodium triacetoxyborohydride (765 mg, 3.63 mmol) was added. The reaction mixture was stirred at room temperature for 5 h. The reaction mixture was partitioned between water (50 mL) and ethyl acetate (25 mL), and the aqueous layer was extracted with ethyl acetate (50 mL x 3). Combined organic phases were washed once with cold water, dried over anhydrous sodium sulfate and the solvent was removed under reduced pressure to furnish the title compound (180 mg, 60% yield), which was used without purification. LC-MS m/z 249.10 [M+H]^+^.

### 3-(4-Chlorophenyl)-1-(4-fluorobenzyl)-1-((4-isopropyl-4*H*-1,2,4-triazol-3-yl)methyl)urea GEN-1

To a stirred solution of *N*-(4-fluorobenzyl)-1-(4-isopropyl-4*H*-1,2,4-triazol-3-yl)methanamine (150 mg, 0.604 mmol) in tetrahydrofuran (10 mL) were added 4-chloroaniline (76 mg, 0.60 mmol) followed by *N,N*-diisopropylethylamine (0.4 mL, 2.41 mmol). Then reaction was cooled to 0 °C, and triphosgene (89 mg, 0.30 mmol) was added. The reaction mixture was stirred at room temperature for 16 h. The reaction mixture was partitioned between water (50 mL) and ethyl acetate (25 mL), and the aqueous layer was extracted with ethyl acetate (25 mL x 3). Combined organic phases were washed once with cold water, dried over anhydrous sodium sulfate and the solvent was removed under reduced pressure to furnish the desired product in its crude form. The crude material was further purified by preparative HPLC (Column: YMC (C18, 20mm X 150mm), Phase A: 0.1% formic acid in water, Phase B: MeCN, Flow: 18ml/min), followed by lyophilization of the fractions, to furnish the title compound as an off-white solid (82.1 mg mg, 33% yield). ^1^H NMR (DMSO-*d*_6_, 400 MHz) *δ* 8.90 (s, 1H), 8.68 (s, 1H), 7.49 (d, *J* = 8.8 Hz, 2H), 7.32-7.28 (m, 4H), 7.19 (t, *J* = 9.2 Hz, 2H), 4.68 (s, 2H), 4.61 (s, 2H), 4.50-4.41 (m, 1H), 1.34 (d, *J* = 6.8 Hz, 6H); LC-MS m/z 402.30 [M+H]^+^.

### Synthesis of 1-(4-Chlorobenzyl)-3-(4-fluorophenyl)-1-((4-isopropyl-4*H*-1,2,4-triazol-3-yl)methyl)urea GEN-2

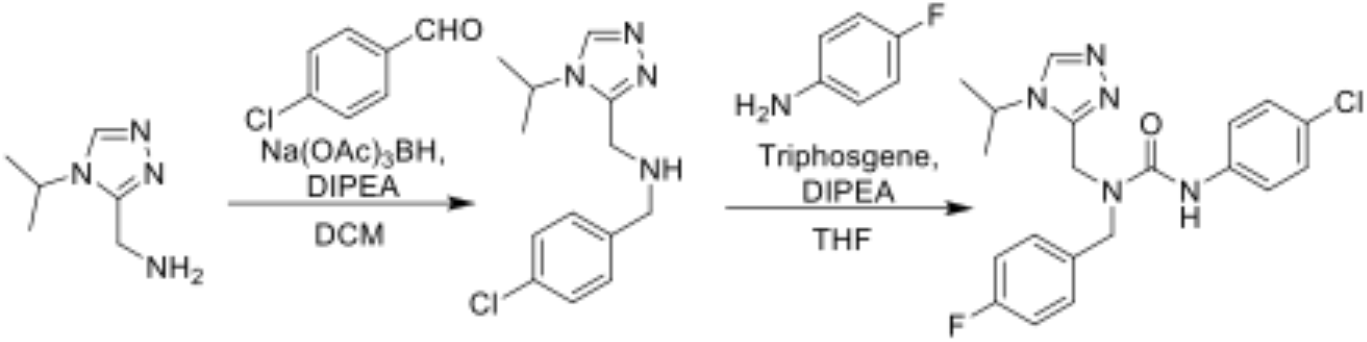

### *N*-(4-Chlorobenzyl)-1-(4-isopropyl-4*H*-1,2,4-triazol-3-yl)methanamine

To a stirred solution of 4-chlorobenzaldehyde (150 mg, 1.07 mmol) in dichloromethane (10 mL) were added (4-isopropyl-4H-1,2,4-triazol-3-yl)methanamine (150 mg, 1.07 mmol), followed by *N,N-* diisopropylethylamine (0.5 mL, 3.214 mmol) and 2 drops of acetic acid. The reaction was cooled to 0 °C, and sodium triacetoxyborohydride (678 mg, 3.21 mmol) was added. The reaction mixture was stirred at room temperature for 5 h. The reaction mixture was partitioned between water (50 mL) and ethyl acetate (25 mL), and the aqueous layer was extracted with ethyl acetate (50 mL x 3). Combined organic phases were washed once with cold water, dried over anhydrous sodium sulfate and the solvent was removed under reduced pressure to furnish the title compound (150 mg, 53% yield), which was used without purification. LC-MS m/z 265.20 [M+H]^+^.

### 1-(4-Chlorobenzyl)-3-(4-fluorophenyl)-1-((4-isopropyl-4*H*-1,2,4-triazol-3-yl)methyl)urea GEN-2

To a stirred solution of *N*-(4-chlorobenzyl)-1-(4-isopropyl-4*H*-1,2,4-triazol-3-yl)methanamine (170 mg, 0.64 mmol) in tetrahydrofuran (10 mL) were added 4-fluoroaniline (71 mg, 0.64 mmol) followed by *N,N-* diisopropylethylamine (0.4 mL, 2.57 mmol). Then reaction was cooled to 0 °C, and triphosgene (95 mg, 0.321 mmol) was added. The reaction mixture was stirred at room temperature for 16 h. The reaction mixture was partitioned between water (50 mL) and ethyl acetate (25 mL), and the aqueous layer was extracted with ethyl acetate (25 mL x 3). Combined organic phases were washed once with cold water, dried over anhydrous sodium sulfate and the solvent was removed under reduced pressure to furnish the desired product in its crude form. The crude material was further purified by preparative HPLC (Column: KINETEX (C18, 21.2mm X 150mm), Phase A: 0.1% TFA in water, Phase B: MeCN, Flow: 18ml/min), followed by lyophilization of the fractions, to furnish the title compound as an off-white solid (18 mg, 6.9% yield). ^1^H NMR (DMSO-*d*_6_, 400 MHz) *δ* 8.80 (s, 1H), 8.68 (s, 1H), 7.45-7.41 (m, 4H), 7.27 (d, *J* = 8.8 Hz, 2H), 7.09 (t, *J* = 8.8 Hz, 2H), 4.68 (s, 2H), 4.61 (s, 2H), 4.51-4.42 (m, 1H), 1.34 (d, *J* = 6.4 Hz, 6H); LC-MS m/z 402.30 [M+H]^+^.

### *T. cruzi* and 3T3 assay protocols

NIH3T3 cells were grown in cell maintenance media consisting of RPMI 1640 supplemented with 1% Penicillin/Streptomycin and 10% fetal bovine serum. *Trypanosoma cruzi* (*T. cruzi*) Tulahuen strain constitutively expressing beta galactosidase activity was propagated by infecting 3T3 cells and collecting supernatants containing trypomastigotes between 5-6 days after NIH3T3 cell infection.

For NIH3T3 cytotox assay, 2000 NIH3T3 cells in 50ml cell maintenance media in the 384 well plates. On the next day, compounds were spotted onto the plates and further incubated for another 4 days at 37°C < 5% CO_2_ incubator. ATP-glo (Promega) was used to evaluate the cytotoxicity of the compound.

For *T. cruzi* assay, 1200 3T3 cells in 40ml of *T. cruzi* assay media (RPMI 1640 with no phenol red, supplemented with 1% Penicillin/Streptomycin and 3% fetal bovine serum) was dispensed in 384 well plate and incubated overnight. On the next day, 5000 *T. cruzi* parasites in 10ml cell maintenance media was introduced and incubated for 1 hour at room temperature. Compounds at various concentrations were spotted and the plates were further Incubated for 6 days at 37°C & 5% CO_2_ incubator. Beta-galactosidase activity was used as a surrogate for presence of parasite10ul of CPRG reagent at final concentration of 100 mM. The plates were incubated for at least 2 hours at room temperature and absorbance measured at 570nm.

## References

1 Gupta, R., Srivastava, D., Sahu, M. et al. Artificial intelligence to deep learning: machine intelligence approach for drug discovery. Mol Divers (2021). https://doi.org/10.1007/s11030-021-10217-3

2 Zhavoronkov, A., Ivanenkov, Y.A., Aliper, A. et al. Deep learning enables rapid identification of potent DDR1 kinase inhibitors. Nat Biotechnol 37, 1038–1040 (2019). https://doi.org/10.1038/s41587-019-0224-x

3 Bian, Y., Xie, XQ. Generative chemistry: drug discovery with deep learning generative models. J Mol Model 27, 71 (2021). https://doi.org/10.1007/s00894-021-04674-8

4 Blaschke, T., Engkvist, O., Bajorath, J. et al. Memory-assisted reinforcement learning for diverse molecular de novo design. J Cheminform 12, 68 (2020). https://doi.org/10.1186/s13321-020-00473-0

5 Popova, M., Isayev, O., & Tropsha, A. (2018). Deep reinforcement learning for de novo drug design. Science advances, 4(7), eaap7885.

6 Automatic Chemical Design Using a Data-Driven Continuous Representation of Molecules Rafael Gómez-Bombarelli, Jennifer N. Wei, David Duvenaud, José Miguel Hernández-Lobato, Benjamín Sánchez-Lengeling, Dennis Sheberla, Jorge Aguilera-Iparraguirre, Timothy D. Hirzel, Ryan P. Adams, and Alán Aspuru-Guzik ACS Central Science 2018 4 (2), 268–276 DOI: 10.1021/acscentsci.7b00572

7 Lim, J., Ryu, S., Kim, J.W. et al. Molecular generative model based on conditional variational autoencoder for de novo molecular design. J Cheminform 10, 31 (2018). https://doi.org/10.1186/s13321-018-0286-7

8 Maziarka, Ł., Pocha, A., Kaczmarczyk, J. et al. Mol-CycleGAN: a generative model for molecular optimization. J Cheminform 12, 2 (2020). https://doi.org/10.1186/s13321-019-0404-1

9 Blanchard, A.E., Stanley, C. & Bhowmik, D. Using GANs with adaptive training data to search for new molecules. J Cheminform 13, 14 (2021). https://doi.org/10.1186/s13321-021-00494-3

10 Méndez-Lucio, O., Baillif, B., Clevert, DA. et al. De novo generation of hit-like molecules from gene expression signatures using artificial intelligence. Nat Commun 11, 10 (2020). https://doi.org/10.1038/s41467-019-13807-w

11 Grechishnikova, D. Transformer neural network for protein-specific de novo drug generation as a machine translation problem. Sci Rep 11, 321 (2021). https://doi.org/10.1038/s41598-020-79682-4

12 All-Assay-Max2 pQSAR: Activity Predictions as Accurate as Four-Concentration IC50s for 8558 Novartis Assays Eric J. Martin, Valery R. Polyakov, Xiang-Wei Zhu, Li Tian, Prasenjit Mukherjee, and Xin Liu Journal of Chemical Information and Modeling 2019 59 (10), 4450–4459 DOI: 10.1021/acs.jcim.9b00375

13 Chatelain E. Chagas disease research and development: Is there light at the end of the tunnel? Comput Struct Biotechnol J. 2016 Dec 14;15:98–103. doi: 10.1016/j.csbj.2016.12.002. PMID: 28066534; PMCID: PMC5196238.

14 Goodfellow, I., Pouget-Abadie, J., Mirza, M., Xu, B., Warde-Farley, D., Ozair, S., … & Bengio, Y. (2014). Generative adversarial nets. Advances in neural information processing systems, 27.

15 Godinez W, Ma E, Chao A, Pei L, Skewes-Cox P, Canham S, et al. JAEGER – Hunting for Antimalarials with Generative Chemistry. ChemRxiv. Cambridge: Cambridge Open Engage; 2021;

16 Jin, Wengong, Regina Barzilay, and Tommi Jaakkola. “Junction tree variational autoencoder for molecular graph generation.” International conference on machine learning. PMLR, 2018.

17 The Synthesizability of Molecules Proposed by Generative Models Wenhao Gao and Connor W. Coley Journal of Chemical Information and Modeling 2020 60 (12), 5714–5723 DOI: 10.1021/acs.jcim.0c00174

18 Ertl, P., Schuffenhauer, A. Estimation of synthetic accessibility score of drug-like molecules based on molecular complexity and fragment contributions. J Cheminform 1, 8 (2009). https://doi.org/10.1186/1758-2946-1-8

19 Bickerton, G., Paolini, G., Besnard, J. et al. Quantifying the chemical beauty of drugs. Nature Chem 4, 90–98 (2012). https://doi.org/10.1038/nchem.1243

20 Rogers, D.; Hahn, M. “Extended-Connectivity Fingerprints.” J. Chem. Inf. and Model. 50:742–54 (2010).

21 Wang, S., & Jiang, J. (2016). A compare-aggregate model for matching text sequences. arXiv preprint arXiv:1611.01747.

22 Salimans, T., Goodfellow, I., Zaremba, W., Cheung, V., Radford, A., & Chen, X. (2016). Improved techniques for training gans. Advances in neural information processing systems, 29, 2234–2242.

23 Abadi, Martín, et al. “Tensorflow: Large-scale machine learning on heterogeneous distributed systems.” arXiv preprint arXiv:1603.04467 (2016).

